# Multisite phosphorylation by Cdk1 initiates delayed negative feedback to control mitotic transcription

**DOI:** 10.1101/2021.02.14.431158

**Authors:** Jonathan B. Asfaha, Mihkel Örd, Christopher R. Carlson, Ilona Faustova, Mart Loog, David O. Morgan

## Abstract

Cell-cycle progression is driven by the phosphorylation of cyclin-dependent kinase (Cdk) substrates^1–3^. The order of substrate phosphorylation depends in part on the general rise in Cdk activity during the cell cycle^4–7^, together with variations in substrate docking to sites on associated cyclin and Cks subunits^3, 6, 8–10^. Many substrates are modified at multiple sites to provide more complex regulation^9, 11–14^. Here, we describe an elegant regulatory circuit based on multisite phosphorylation of Ndd1, a transcriptional co-activator of genes required for mitotic progression^15, 16^. As cells enter mitosis, Ndd1 phosphorylation by Cdk1 is known to promote mitotic cyclin (*CLB2*) gene transcription, resulting in positive feedback^17–20^. Consistent with these findings, we show that low Cdk1 activity promotes *CLB2* expression at mitotic entry. We also find, however, that *CLB2* expression is inhibited by high levels of Cdk1 activity in a mitotic arrest. Inhibition is accompanied by Ndd1 degradation, and we present evidence that high mitotic Cdk1-Clb2 activity generates phosphodegrons on Ndd1, leading to its degradation. Complete Ndd1 phosphorylation by the Clb2-Cdk1-Cks1 complex requires the phosphothreonine-binding site of Cks1, as well as a novel phosphate-binding pocket on the cyclin Clb2^21^. We therefore propose that initial phosphorylation by Cdk1 primes the protein for secondary phosphorylation at phosphodegrons, resulting in degradation only at high Cdk1 activity. Together, our results suggest that rising levels of mitotic Cdk1 activity act at multiple phosphorylation sites on Ndd1, first triggering rapid positive feedback and then promoting delayed negative feedback, resulting in a pulse of mitotic gene expression.

## Results

### Negative feedback suppresses mitotic gene expression in a mitotic arrest

We analyzed expression of the *CLB2* cluster of mitotic genes^16^ by measuring mRNA levels with RT-qPCR. As in previous studies, we found that expression of genes in this cluster reaches peak levels in mitosis, 60 min after release from a G1 arrest (Figure 1A)^16^. Transcriptional activation is driven by M-phase Cdk1 activity^4, 22, 23^. We therefore expected that high Cdk1 activity in a mitotic arrest would sustain continuous transcription of mitotic genes. To test this possibility, we arrested cells in G1 and released them into nocodazole, a microtubule poison that activates the spindle assembly checkpoint and arrests cells in mitosis. Surprisingly, we found that upon entry into a mitotic arrest, expression of mitotic genes decreased to between 50% and 20% of peak levels (Figure 1B). Similar results were obtained when cells were arrested in mitosis by *CDC20* depletion, indicating that the effect was not due to spindle assembly checkpoint activation (Figure 1C).

**Figure 1.**
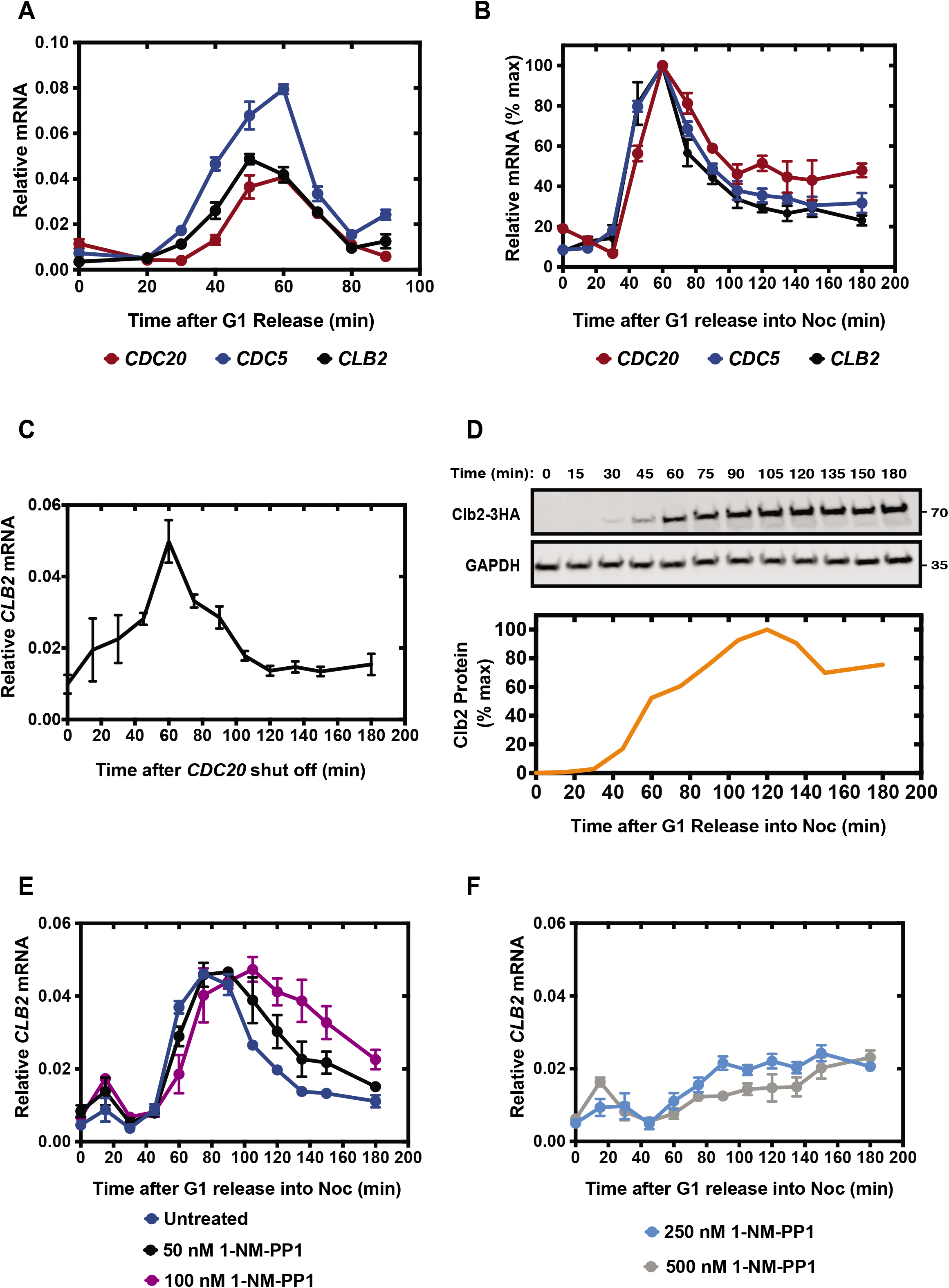
Decline in *CLB2* cluster gene expression in a mitotic arrest depends on Cdk1 activity. (A) Cells were released from a G1 arrest and harvested at the indicated times for analysis of gene expression by RT-qPCR. mRNA for each of the genes was normalized to *ACT1* mRNA. Data points indicate mean +/-S.E.M. in 3 independent experiments. (B) Gene expression was measured in cells released from G1 arrest into nocodazole (Noc) to induce a mitotic arrest. mRNA for each of the genes was normalized to *ACT1* mRNA. Data points indicate mean +/-S.E.M. in 3 independent experiments. (C) Cells carrying an endogenous replacement of the *CDC20* promoter with a galactose-inducible promoter were released from a G1 arrest into dextrose to induce a mitotic arrest. *CLB2* expression was measured at the indicated times by RT-qPCR (normalized to *ACT1* mRNA). Data points indicate mean +/-S.E.M. in 3 independent experiments. (D) Cells were released from G1 into nocodazole. Expression of 3xHA-tagged Clb2 was analyzed by immunoblot at the indicated times, with a GAPDH immunoblot serving as a loading control. (E & F) Cells carrying analog-sensitive Cdk1 (*cdk1-as1*) were released from G1 arrest into media containing the indicated concentrations of 1-NM-PP1, plus nocodazole. *CLB2* gene expression was measured at the indicated times by RT-qPCR (normalized to *ACT1* mRNA). Data points indicate mean +/-S.E.M. in 3 independent experiments.

Despite the reduction in transcript levels, Clb2 protein levels remained at a constant high level in a mitotic arrest, as seen in previous studies (Figure 1D)^24–26^. Clb2 is very stable in a mitotic arrest, which would explain its high steady-state levels despite the decline in its mRNA transcript.

Cdk1-mediated positive feedback promotes *CLB2* transcriptional activation in mitosis^18, 19, 23, 27^. We hypothesized that high mitotic Cdk1 activity might suppress *CLB2* transcription. We inhibited Cdk1 at various inhibitor concentrations in an analog-sensitive Cdk1 mutant, *cdk1-as1*. Cdk1 in this strain bears an F88G mutation in the active site, making Cdk1 highly selective for the inhibitor 1-NM-PP1^28^. Mild Cdk1 inhibition, at 50 nM or 100 nM 1-NM-PP1, delayed the reduction in *CLB2* transcription (Figure 1E). Mild inhibition did not significantly affect the initial mitotic rise in *CLB2* expression, suggesting that partial Cdk1 inhibition did not affect transcriptional activation. However, strong Cdk1 inhibition, at 250 nM or 500 nM 1-NM-PP1, suppressed transcription (Figure 1F). Thus, strong Cdk1 inhibition inhibits positive feedback and transcriptional activation, while mild inhibition blocks negative feedback while leaving positive feedback less affected. A simple explanation for these results is that negative feedback requires higher levels of kinase activity and is therefore more readily inhibited by partial kinase inhibition.

### Ndd1 is destabilized in a mitotic arrest by N-terminal phosphorylation sites

Ndd1 is a key positive regulator of the *CLB2* transcriptional cluster and is activated by Cdk1-mediated phosphorylation^19^. We therefore tested its role in the negative feedback we observed. Immunoblotting revealed that Ndd1 levels decreased as cells proceeded into a mitotic arrest from G1, and this decline could be alleviated by the addition of 100 nM 1-NM-PP1 (Figure 2A, Figure S1A).

**Figure 2.**
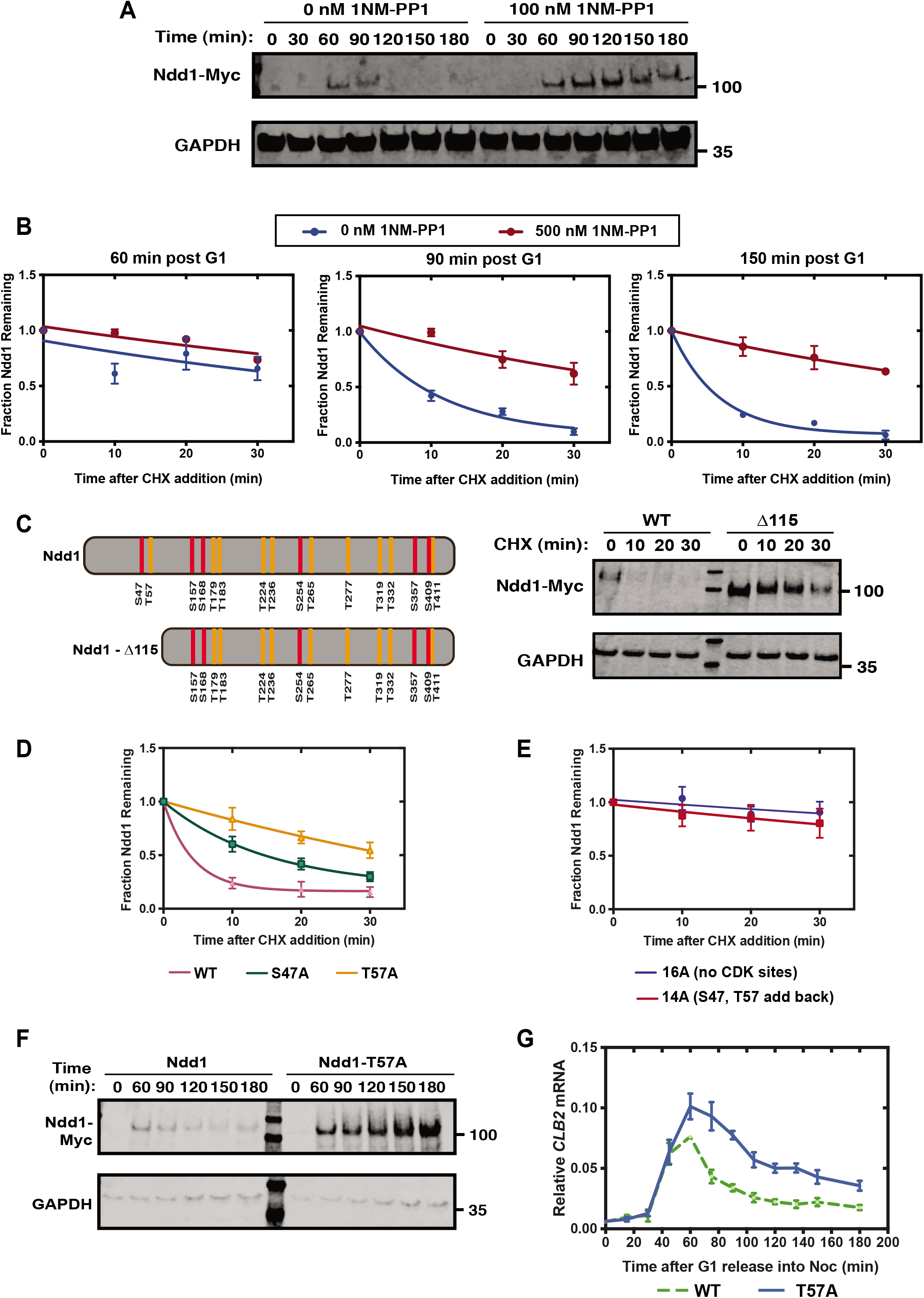
Ndd1 destabilization depends on rising Cdk1 activity and N-terminal phosphorylation. (A) c*dk1-as1* cells were released from a G1 arrest into nocodazole in the absence (left) or presence (right) of 100 nM 1-NM-PP1. Myc-tagged Ndd1 and GAPDH expression were monitored by immunoblot at the indicated times. (B) Parallel cultures of *cdk1-as1* cells were released from a G1 arrest into nocodazole, in the presence or absence of 500 nM 1-NM-PP1. Cycloheximide (CHX) was added after 60, 90, or 150 min. Samples were harvested after 0, 10, 20, or 30 min in CHX. Myc-tagged Ndd1 levels were analyzed by immunoblot and quantified. Data points indicate mean +/-S.E.M. in 2 independent experiments. Curves were generated by fitting data to a single exponential decay model. (C) *Left:* Depiction of Ndd1 and an Ndd1 mutant lacking the first 115 amino acids (Ndd1-Δ115), with 16 consensus Cdk1 sites marked (serine, red; threonine, orange). *Right:* Cells carrying galactose-inducible wild-type (WT) Ndd1 or Ndd1-Δ115 were grown in galactose and arrested in mitosis with nocodazole. CHX was added, and Ndd1-Myc and GAPDH were analyzed by immunoblotting at the indicated times. (D) Cells carrying Ndd1 (WT), Ndd1-S47A, or Ndd1-T57A were treated as in panel C. Quantification of immunoblots from 3 independent experiments (mean +/- S.E.M.). Curves were generated by fitting data to a single exponential decay model. Representative immunoblot in Figure S1B. (E) Cells carrying Ndd1-16A or Ndd1 containing only S47 and T57 (Ndd1-14A) were treated as in panel C. Quantification of immunoblots from 3 independent experiments (mean +/- S.E.M.). Curves were generated by fitting data to a single exponential decay model. Representative immunoblot in Figure S1C. (F) Cells carrying Myc-tagged WT Ndd1 or Ndd1-T57A at the endogenous *NDD1* locus were released from a G1 arrest into nocodazole to induce a mitotic arrest. Myc-tagged Ndd1 and GAPDH levels were analyzed by immunoblotting at the indicated times. (G) Cells carrying WT Ndd1 or Ndd1-T57A as in panel F were released from G1 arrest into nocodazole. Samples were harvested at the indicated times for analysis of *CLB2* gene expression by RT-qPCR (normalized to *ACT1* mRNA). Dashed line indicates data for WT from Figure 1B. Data points indicate mean +/-S.E.M. in 3 independent experiments.

We measured the effect of Cdk1 inhibition on Ndd1 stability at various points after release from G1. We estimated Ndd1 half-life by measuring its decline after addition of cycloheximide at 60, 90, or 150 min after release from G1 into a mitotic arrest. We found that Ndd1 degradation rate increased as cells proceeded into the mitotic arrest (Figure 2B; blue). Inhibition of Cdk1 with 500 nM 1-NM-PP1 prevented Ndd1 degradation at all time points (Figure 2B; red). These data indicate that Cdk1-dependent Ndd1 degradation rate increases gradually during a mitotic arrest, suggesting that prolonged high Cdk1 activity promotes degradation.

Ndd1 is reminiscent of the many other proteins that are destabilized by Cdk1-mediated phosphorylation. In most of these cases, Cdk1 phosphorylation sites generate phosphodegron sequence motifs recognized by F-box subunits of the ubiquitin ligase SCF^11, 29–31^. Interestingly, the F-box protein Grr1 is known to be required for Ndd1 degradation in a spindle checkpoint arrest^32^, suggesting that phosphorylated Ndd1 is ubiquitylated by the ubiquitin ligase SCF^Grr1^, leading to its proteasomal destruction.

Ndd1 contains 16 Cdk1 consensus phosphorylation sites (minimal site: S/T-P; full site: S/T-P-x-K/R), most of which are conserved in budding yeasts (Figure 2C, Figure S2). Mutation of all 16 sites is known to stabilize Ndd1 in a spindle checkpoint arrest^32^. To identify the specific sites in Ndd1 that are required for degradation, we first assessed the stability of Ndd1 truncation mutants. Deletion of the amino-terminal 115 amino acids stabilized Ndd1 (Figure 2C). Two minimal Cdk1 consensus sites, S47 and T57, are found in this region. Consistent with the importance of these sites, an S47A substitution partially stabilized Ndd1, while a T57A substitution led to a greater level of stabilization (Figure 2D, Figure S1B).

Mutation of all 16 Cdk1 sites (the Ndd1-16A mutant) resulted in greater Ndd1 stabilization than that seen with the T57A mutant (Figure 2E, blue; Figure S1C). Adding back S47 and T57 to the 16A mutant did not induce degradation (Figure 2E, red; Figure S1C). Thus, S47 and T57 are required but not sufficient for rapid Ndd1 degradation.

Ndd1 is required for high mitotic *CLB2* transcription^19^, and therefore Ndd1 degradation is likely to be responsible, at least in part, for the decline in *CLB2* transcription during a mitotic arrest. We tested the importance of Ndd1 degradation by analyzing *CLB2* mRNA levels in a yeast strain in which *NDD1* was replaced with a mutant gene encoding Ndd1-T57A. When cells expressing this mutant protein were released from a G1 arrest into nocodazole, Ndd1-T57A protein levels remained high (Figure 2F). This mutant did not affect the initial rate of increase in *CLB2* transcription or the timing of the peak of maximum transcription. *CLB2* mRNA levels accumulated to higher levels in the mutant strain (Figure 2G), but a gradual decline in *CLB2* mRNA was still apparent. Thus, although Ndd1 degradation is likely to contribute to decreased *CLB2* transcription, additional mechanisms must inhibit Ndd1 or other transcriptional regulators.

### Ndd1 degradation depends on multiple Clb2-dependent phosphorylation sites

Because S47 and T57 are not sufficient to induce degradation, and their mutation does not fully stabilize Ndd1, we searched for other sites that contribute to Ndd1 destabilization. We systematically added back clusters of predicted Cdk1 sites to Ndd1-16A, starting at the amino terminus, and measured the stability of each mutant (Figure 3A). Adding back predicted Cdk1 sites up to and including T236 (the Ndd1-8A mutant) had minimal effects on degradation rate (Figure 3B). However, further addition of S254 and T265 (the Ndd1-6A mutant) led to a decline in stability (Figure 3C). Addition of more predicted C-terminal Cdk1 sites further destabilized the protein (Figure 3C).

**Figure 3.**
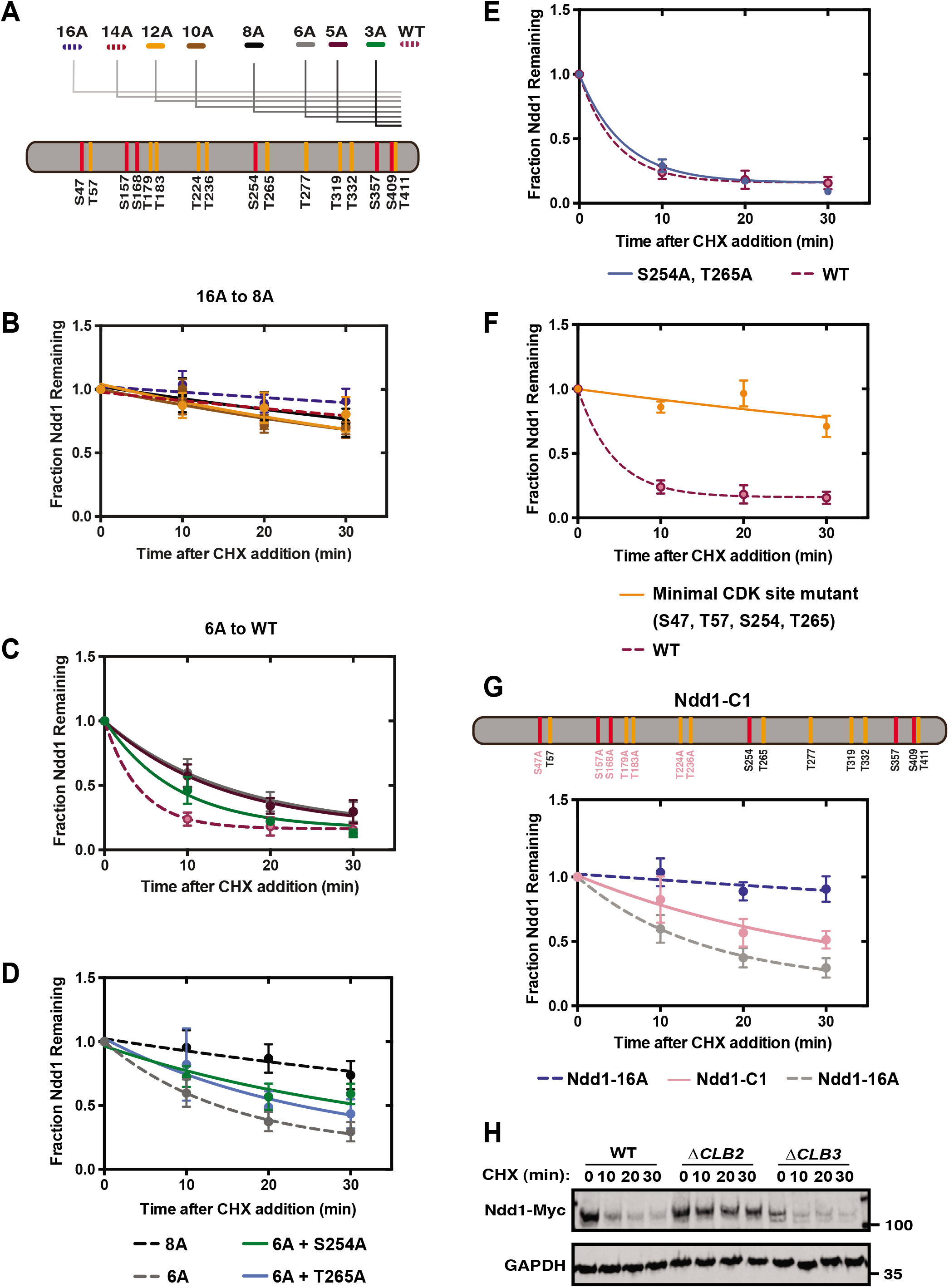
Multiple predicted Cdk1 sites promote Ndd1 destabilization. (A) Diagram of predicted Cdk1 sites in Ndd1, showing the sites mutated to alanine in the Ndd1 mutants analyzed in panels B and C. (B) Cells carrying galactose-inducible Ndd1 alanine mutants 12A, 10A, or 8A (colors indicated in panel A) were grown in galactose and arrested in mitosis with nocodazole. CHX was added, and Ndd1-Myc was quantified by immunoblotting at the indicated times. Data points indicate mean +/-S.E.M. in 3 independent experiments. Dashed lines indicate data for Ndd1-16A and Ndd1-14A from Figure 2E. (C) Cells carrying galactose-inducible Ndd1 alanine mutants 6A, 5A, or 3A were treated and analyzed as in panel B. Data points indicate mean +/-S.E.M. in 3 independent experiments. Dashed line indicates data for WT Ndd1 from Figure 2D. (D) Cells were constructed in which galactose-inducible Ndd1 carried alanine mutations at the C-terminal six Cdk1 consensus sites (Ndd1-6A; see panel A) plus a mutation of either S254 or T265 to alanine, and were treated as in panel B. Quantification of immunoblots from 3 independent experiments (mean +/- S.E.M.). Dashed lines indicate data for Ndd1-8A and Ndd1-6A from panels B and C, respectively. Representative immunoblot in Figure S3A. (E) Cells carrying galactose-inducible Ndd1-S254A, T265A were treated and analyzed as in panel B. Quantification of immunoblots from 3 independent experiments (mean +/- S.E.M.). Dashed line indicates data for WT Ndd1 from Figure 2D. Representative immunoblot in Figure S3B. (F) Cells were constructed in which galactose-inducible Ndd1 carries alanine mutations at all Cdk1 consensus sites except S47, T57, S254, and T265. Cells were treated and analyzed as in panel B. Quantification of immunoblots from 3 independent experiments (mean +/- S.E.M.). Dashed line indicates data for WT Ndd1 from Figure 2D. Representative immunoblot in Figure S3C. (G) Cells carrying galactose-inducible Ndd1 containing only T57, S254, T265, T277, T319, T332, S357, S409, and T411 (Ndd1-C1) were treated and analyzed as in panel B. Quantification of immunoblots from 3 independent experiments (mean +/- S.E.M.). Dashed lines indicate data for Ndd1-16A and Ndd1-6A from Figure 2E and panel C, respectively. Representative immunoblot in Figure S3D. (H) Cells expressing Ndd1-Myc under the control of the *GAL* promoter were grown in galactose and arrested in mitosis with nocodazole. CHX was added, and Ndd1-Myc and GAPDH were analyzed by immunoblotting at the indicated times.

We added back either S254 or T265 to Ndd1-8A. Addition of either site led to an intermediate level of destabilization that fell between those of Ndd1-8A and Ndd1-6A (Figure 3D, Figure S3A). Thus, both sites promote instability. Interestingly, however, when S254 and T265 are both mutated to alanine in an otherwise wild-type protein, Ndd1 is not stabilized (Figure 3E, Figure S3B). Furthermore, a version of Ndd1 that contains only S47, T57, S254, and T265 is stable (Figure 3F, Figure S3C), showing that these four sites, despite their contributions in some contexts, are not sufficient for degradation. Finally, a mutant containing only T57, S254, T265, and all sites C-terminal of T265, was partially destabilized (Figure 3G, Figure S3D), consistent with evidence in Figure 4C that sites C-terminal of T265 are sufficient to promote some degradation. Together, our studies of numerous site mutations suggest that multiple sites collaborate to induce Ndd1 degradation.

**Figure 4.**
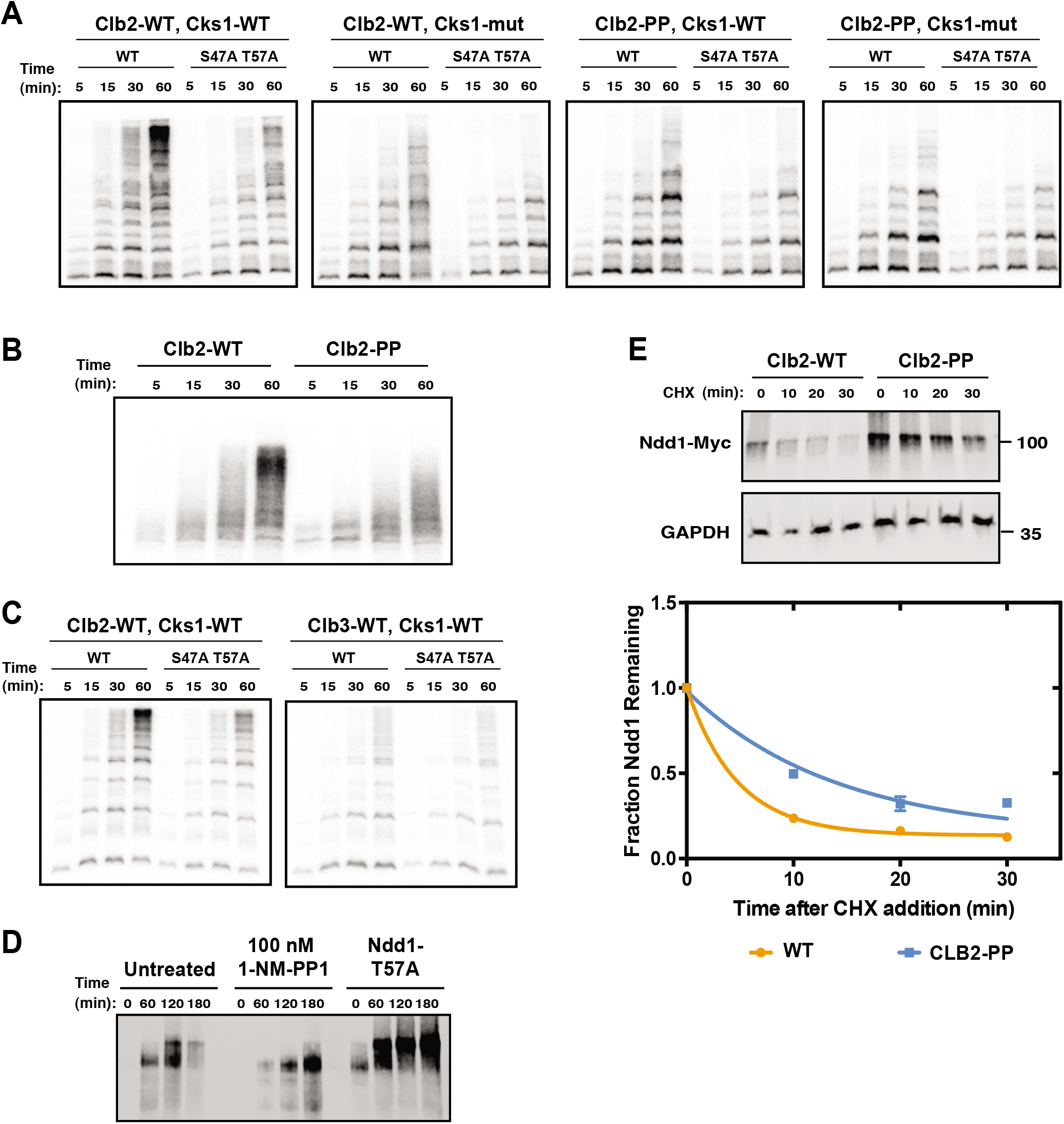
N-terminal phosphorylation of Ndd1 depends on phosphate-binding pockets of Cks1 and Clb2. (A) Truncated Ndd1 (aa 1-272), either wild-type (WT) or S47A T57A, was incubated with [γ-^32^P]-ATP and the indicated Clb2-Cdk1-Cks1 complex: panel 1: wild-type complex; panel 2: a complex containing Cks1 with mutations in the phosphothreonine-binding site (R33E S82E R102A; Cks1-mut); panel 3: a complex containing Clb2 with mutations in the phosphate-binding pocket (R366A R379A K383A; Clb2-PP); panel 4: a complex containing mutations in both Cks1 and Clb2. Reactions were stopped at the indicated times, and reaction products were analyzed by Phos-tag SDS-PAGE and autoradiography. Results are representative of three independent experiments. (B) Full-length Ndd1-WT was incubated with [γ-^32^P]-ATP and wild-type (WT) Clb2-Cdk1-Cks1 or with a complex containing Clb2-PP. Reactions were analyzed as in panel A. Results are representative of two independent experiments. (C) Truncated Ndd1 (aa 1-272), either wild-type (WT) or S47A T57A, was incubated with [γ-^32^P]-ATP and either Clb2-Cdk1-Cks1 (left) or Clb3-Cdk1-Cks1 (right). Reactions were analyzed as in panel A. Results are representative of two independent experiments. (D) c*dk1-as1* cells carrying Myc-tagged wild-type Ndd1 (left 8 lanes), or wild-type cells carrying Myc-tagged Ndd1-T57A (right 4 lanes), were released from a G1 arrest into nocodazole. As indicated, some cells were treated with 100 nM 1-NM-PP1 to inhibit Cdk1 activity. Levels and phosphorylation of Ndd1 were assessed by Phos-tag SDS-PAGE and immunoblotting. (E) A strain was constructed in which the *CLB2* locus was replaced with a mutant gene encoding Clb2-PP. Cells carrying galactose-inducible Ndd1-Myc were grown in galactose and arrested in mitosis with nocodazole. CHX was added, and Ndd1-Myc was analyzed by immunoblotting at the indicated times. Top panel shows a representative immunoblot. Bottom panel shows quantification of Ndd1 from 4 independent experiments (mean +/- S.E.M.).

If Ndd1 stability is tied to cell cycle stage, then cyclin specificity may be an important determinant of Ndd1 destruction. The two major mitotic cyclins, Clb3 and Clb2, are thought to act in sequence as cells approach and progress into mitosis, so we tested their contributions to Ndd1 stability in a mitotic arrest. Deletion of Clb3 had no effect, whereas deletion of Clb2 increased the stability of Ndd1 (Figure 3H). Thus, Ndd1 instability depends primarily on high Clb2-dependent Cdk1 activity during a prolonged mitotic arrest.

To gain more insight into the roles of specific sites, we used mass spectrometry to identify major Cdk1-dependent phosphorylation sites in vitro. Previous biochemical studies have demonstrated that Ndd1 is an excellent substrate of both Clb3-Cdk1 and Clb2-Cdk1^12, 33, 34^. We confirmed that purified Clb2-Cdk1-Cks1 complexes phosphorylate multiple sites in Ndd1, with some preference for S254 and other sites in the amino-terminal region (Figure S4A). We also compared the rates of phosphorylation by Clb2-Cdk1-Cks1 and Clb3-Cdk1-Cks1 at specific sites (Figure S4B). We observed moderate Clb3 specificity for sites C-terminal of T277, including T319, which is required for Ndd1 activation^19^. In the amino-terminal half of the protein, T265 displayed moderate Clb2 specificity. Unfortunately, phosphorylation at the two key N-terminal sites, S47 and T57, was not measurable in these experiments because the peptide containing these sites was not detected in the mass spectrometer (Figure S5). We therefore turned to other methods for analysis of these sites, as described next.

### Phosphorylation of N-terminal sites depends on priming by phosphate-binding sites in Cks1 and Clb2

The requirement for multiple phosphorylation sites for Ndd1 degradation is reminiscent of other Cdk1 substrates, including the Cdk1 inhibitor Sic1. In many of these cases, some phosphorylation sites are destabilizing not because they form a phosphodegron but because they act as priming sites to enhance secondary phosphorylation at a phosphodegron. These mechanisms often depend on the phosphothreonine-binding site of the associated Cks1 subunit, which interacts with priming sites to promote phosphorylation at other sites^6, 9, 12, 35^.

With these ideas in mind, we used Phos-tag SDS-PAGE to analyze the stoichiometry of Ndd1 phosphorylation by Cdk1 in vitro^36^. To improve gel resolution, we used a truncated protein containing the N-terminal 272 residues, which contains ten Cdk1 sites, including those most clearly implicated in degradation (S47, T57, S254, T265). Ndd1 is predicted to be disordered, so truncation is unlikely to disrupt a folded domain. Phosphorylation by wild-type Clb2-Cdk1-Cks1 led to phosphorylation of many sites on this protein (Figure 4A; panel 1). Mutation of S47 and T57 greatly reduced the formation of hyperphosphorylated Ndd1, demonstrating that one or both of these sites are substrates of Clb2-Cdk1-Cks1. The S47A T57A mutations had little effect on the hypophosphorylated forms seen at earlier time points in the reaction, indicating that S47 and T57 are phosphorylated at a lower rate than several rapidly phosphorylated initial sites.

Mutation of the phosphothreonine-binding pocket of Cks1 greatly reduced the hyperphosphorylated Ndd1 forms, but had a minor impact on the primary phosphorylation sites at early time points (Figure 4A; panel 2). S47A T57A mutations further reduced the hyperphosphorylated forms but had little effect on rapidly phosphorylated forms. Cks1 therefore enhances phosphorylation of multiple weaker sites, including S47 and T57.

Recent structural studies led to the discovery of a previously uncharacterized phosphate-binding pocket on the surface of human cyclin B1^21^. The three basic residues that form this pocket are highly conserved throughout the eukaryotic B-type cyclins and are found in all six of the budding yeast Clb proteins. We hypothesized that this pocket in Clb2 might act, like Cks1, to enhance phosphorylation of Ndd1 following phosphorylation at priming sites. Consistent with this idea, we found that mutation of the three key residues in the Clb2 pocket (R366A R379A K383A, termed Clb2-PP) reduced Ndd1 phosphorylation, particularly at sites modified at later time points (Figure 4A; panel 3). Mutation of S47 and T57 reduced phosphorylation at these sites further. The Clb2 phosphate-binding pocket was also required for full phosphorylation of full-length Ndd1 (Figure 4B).

When we combined mutations in the phosphate-binding sites of both Cks1 and Clb2, hyperphosphorylation of Ndd1 was almost abolished (Figure 4A; panel 4). Mutation of S47 and T57 in this context had relatively little impact, suggesting that these sites depend largely on priming phosphorylation at other sites.

We also compared the patterns of phosphorylation of Ndd1(1-272) with Clb2 and Clb3. Clb3 was less effective in catalyzing hyperphosphorylation, including S47 and T57, consistent with our evidence that it does not contribute significantly to Ndd1 degradation in the cell (Figure 4C).

Our results raised the possibility that the phosphodegrons of Ndd1, like those in Sic1, contain suboptimal phosphorylation sites: that is, poor sites that are extensively modified only after priming at other sites. S47 and T57 are reasonable candidates for one phosphodegron: these sites are required but not sufficient for Ndd1 degradation, and we observe negligible phosphorylation of a mutant Ndd1 with T57 as the only Cdk1 site (Figure S4A). We obtained further evidence for this model by analyzing the phosphorylation state of Ndd1 in the cell by Phos-tag gels, which showed that Ndd1-T57A accumulates in a highly phosphorylated state despite being highly stabilized (Figure 4D).

Importantly, we also obtained evidence that priming is required for rapid Ndd1 degradation, using a yeast strain in which the *CLB2* gene was replaced with a gene encoding the Clb2-PP mutant. This strain did not display major cell cycle defects, as expected given that *clb2Δ* cells are viable. In nocodazole-arrested cells, steady-state levels of Ndd1 were higher in this strain than in the wild type, and half-life analysis indicated that Ndd1 was partially stabilized in mutant cells (Figure 4E). These results suggest that the phosphate-binding pocket of Clb2 helps promote secondary phosphorylation at sites, perhaps including S47 and T57, that serve as phosphodegrons in vivo.

## Discussion

Expression of genes in the *CLB2* transcriptional cluster is known to be stimulated by Cdk1 activity, suggesting that the mitotic rise in *CLB2* expression depends, at least in part, on positive feedback^18–20, 23, 27^. Our work suggests that early Cdk1 activity at the *CLB2* promoter stimulates the production of Clb2-Cdk1 activity as cells enter mitosis, after which accumulation of high Clb2-Cdk1 activity leads to delayed negative feedback that inhibits transcription, in part through Ndd1 degradation and in part through other, as yet unidentified, mechanisms. This time-delayed negative feedback system generates a pulse of mitotic transcription in a mitotic arrest^37^.

Our studies of the stability of various Ndd1 phosphomutants, together with biochemical analyses of Ndd1 phosphorylation in vitro, suggest that full Ndd1 phosphorylation and degradation depend on sequential cascades of phosphorylation. Rapidly phosphorylated primary sites interact with phosphate-binding sites on Cks1 and Clb2, which leads to delayed, partial secondary phosphorylation at suboptimal sites that promote degradation. Inside the cell, we speculate that these mechanisms create a system in which degradation occurs only when kinase activity rises above a threshold that overcomes opposing phosphatases acting on the same degradative sites^10, 12, 13^.

Our work provides evidence that a recently described phosphate-binding pocket on B-type cyclins^21^ contributes to the regulation of Cdk1 substrate phosphorylation. Cks1 is known to direct phosphorylation downstream of priming phosphothreonine sites^10, 12^, but the specificity of the cyclin phosphate-binding pocket is not known. Detailed further studies with an array of Ndd1 mutants will be required to clarify which sites act as priming sites, and whether Cks1 and Clb2 act in sequence or simultaneously in promoting secondary phosphorylation. Additional studies will also be required to assess the role of the cyclin pocket in phosphorylation of other substrates or regulators. There is evidence, for example, that the residues in this pocket influence interactions between cyclin and the regulatory phosphatase Cdc25^38^.

The structure of Ndd1 phosphodegrons remains poorly understood. Cdk1-dependent Ndd1 degradation is likely to result from the binding of phosphodegrons to the Grr1 F-box subunit of the ubiquitin ligase SCF^32^. Although multiple substrates of Grr1 have been discovered, its mechanisms of substrate interaction are not defined. The central leucine-rich repeat (LRR) domain of Grr1 forms an arc-like structure comprised of an outer arc of alpha helices that positions an inner surface of beta-sheets, which contain multiple positively-charged residues that are necessary for recognizing phosphosites on Grr1-specific substrates. Mutations that span the inner cationic surface of Grr1 disrupt substrate binding, suggesting that Grr1 contains multiple phosphate-binding sites^31^. Regions that destabilize Grr1 substrates, as seen in Cln2 and Hof1, are enriched in clusters of phosphorylation sites that collectively destabilize these proteins^30, 39–41^. Thus, Grr1 might recognize clusters of phosphorylation, as opposed to the relatively well-defined diphosphodegron recognized by the F-box protein Cdc4.

Many of the Cdk1 sites in Ndd1 are grouped into pairs of sites 10-12 residues apart (Figure S2). This spacing is seen for S47/T57 and for S254/T265, which contribute to degradation. Perhaps these pairs serve as weak diphosphodegrons dispersed at multiple locations: it is conceivable, for example, that T57 and S47, or S254 and T265, each encode specificity for Grr1 but do not bind with sufficient affinity to maintain a stable interaction. The avidity imparted by multiple weakly-interacting phosphosites binding to the cationic surface of Grr1 might be required for stable association and degradation.

If later Clb2-dependent phosphorylation events destabilize Ndd1, then early phosphorylation events, catalyzed by Clb3-Cdk1, are likely to initiate the activation of Ndd1. Indeed, previous studies suggest that *CLB2* transcription is partially reduced in the absence of Clb3^23^. Cdk1-dependent phosphorylation of T319 on Ndd1 is critical for its promoter recruitment and thus for *CLB2* transcriptional activation^19^. Our mass spectrometry studies revealed that T319 displays moderate specificity for Clb3 over Clb2 (Figure S4B), and recent evidence suggests that T319 phosphorylation is reduced in a strain with Clb2 as the only B-type cyclin^7^. Other studies have demonstrated that Ndd1 phosphorylation depends in part on the substrate-docking pocket of Clb3^33^. Clb3 might therefore initiate Ndd1 activation before positive feedback is triggered by Clb2 (and Cdc5^42^). Upon entry into mitosis, Clb2-Cdk1 alone is capable of catalyzing phosphorylation of the key degradative sites in Ndd1, perhaps due to its higher intrinsic activity^34^, its higher concentration in the nucleus^10^, or specificity in its phosphate-binding pocket.

Multiple sites in the C-terminal region of Ndd1 enhance the rate of Ndd1 degradation (Figures 3C,G). Some of these sites display moderate Clb3 specificity (Figure S4B), suggesting that early Clb3-dependent C-terminal phosphorylation contributes to Ndd1 degradation as the cell enters mitosis. However, although Clb3 is abundant in nocodazole-arrested cells^26^, our results argue that it is not required for Ndd1 degradation in those cells (Figure 3H). Phosphorylation of C-terminal sites by Clb2-Cdk1 appears to be sufficient to enhance degradation in a mitotic arrest.

Hundreds of Cdk1 substrates are phosphorylated at specific times during progression through the cell cycle^1–3^. Many of these proteins contain arrays of phosphorylation sites in disordered regions. The timing and switch-like behavior of phosphorylation are governed by a remarkably complex variety of parameters, including the sequences, affinities, positioning, and distancing of phosphosites and cyclin-specific docking motifs^9^. Ndd1 appears to provide an intriguing case in which multisite phosphorylation of a single protein, by sequential cyclin-Cdk1 complexes, provides ordered activation and inactivation of a specific cell cycle process.

## STAR METHODS

### KEY RESOURCES TABLE

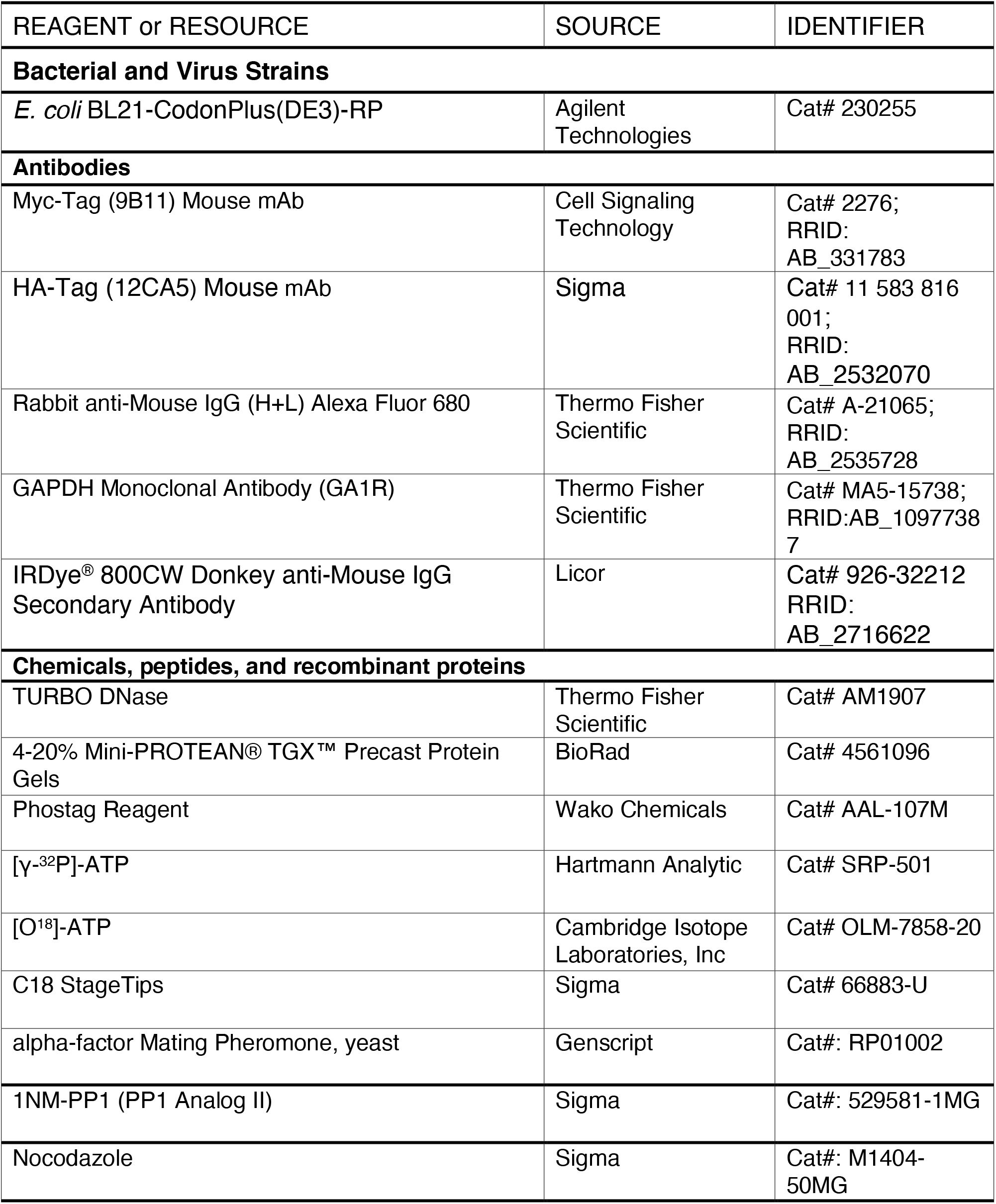
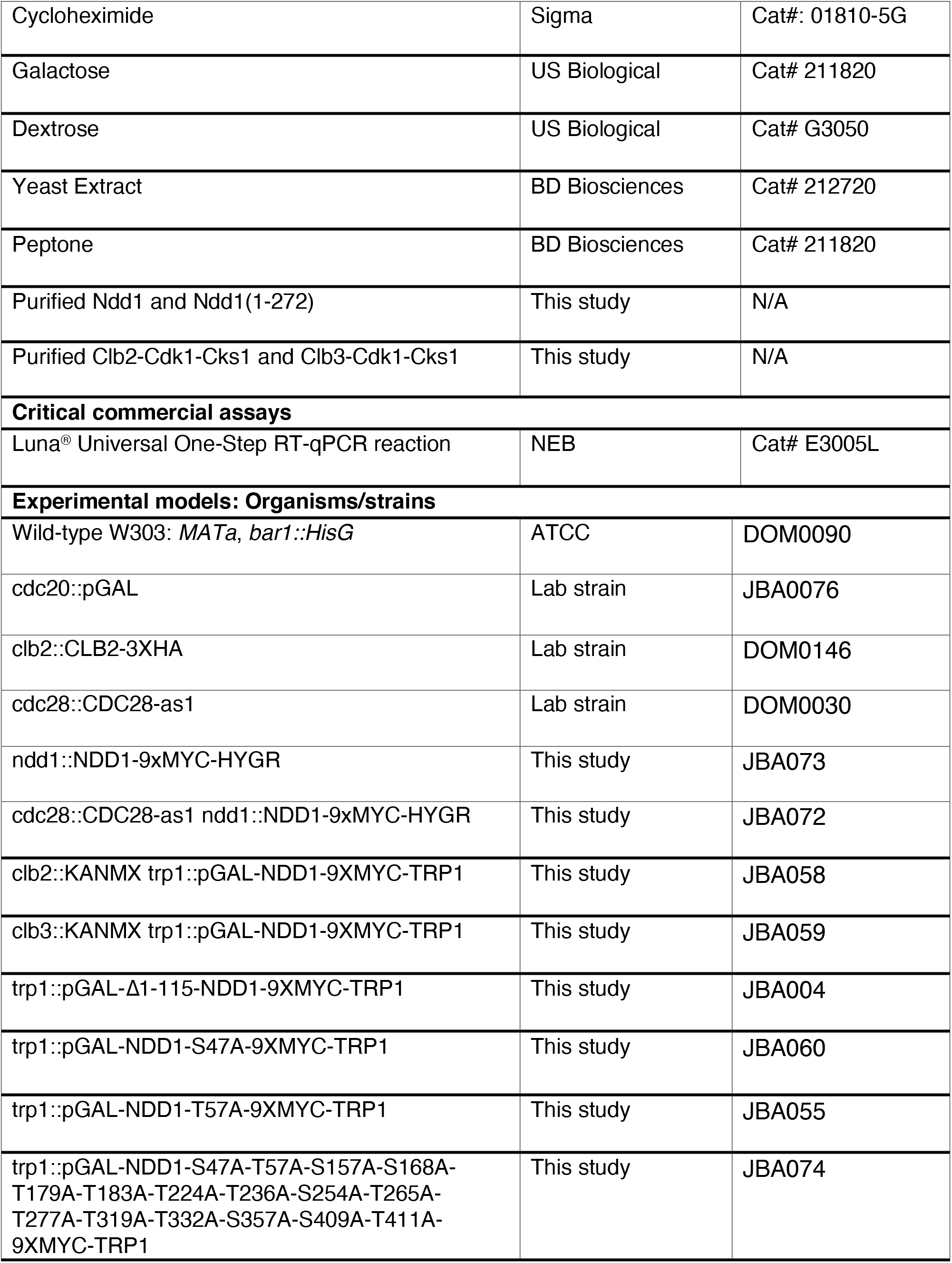
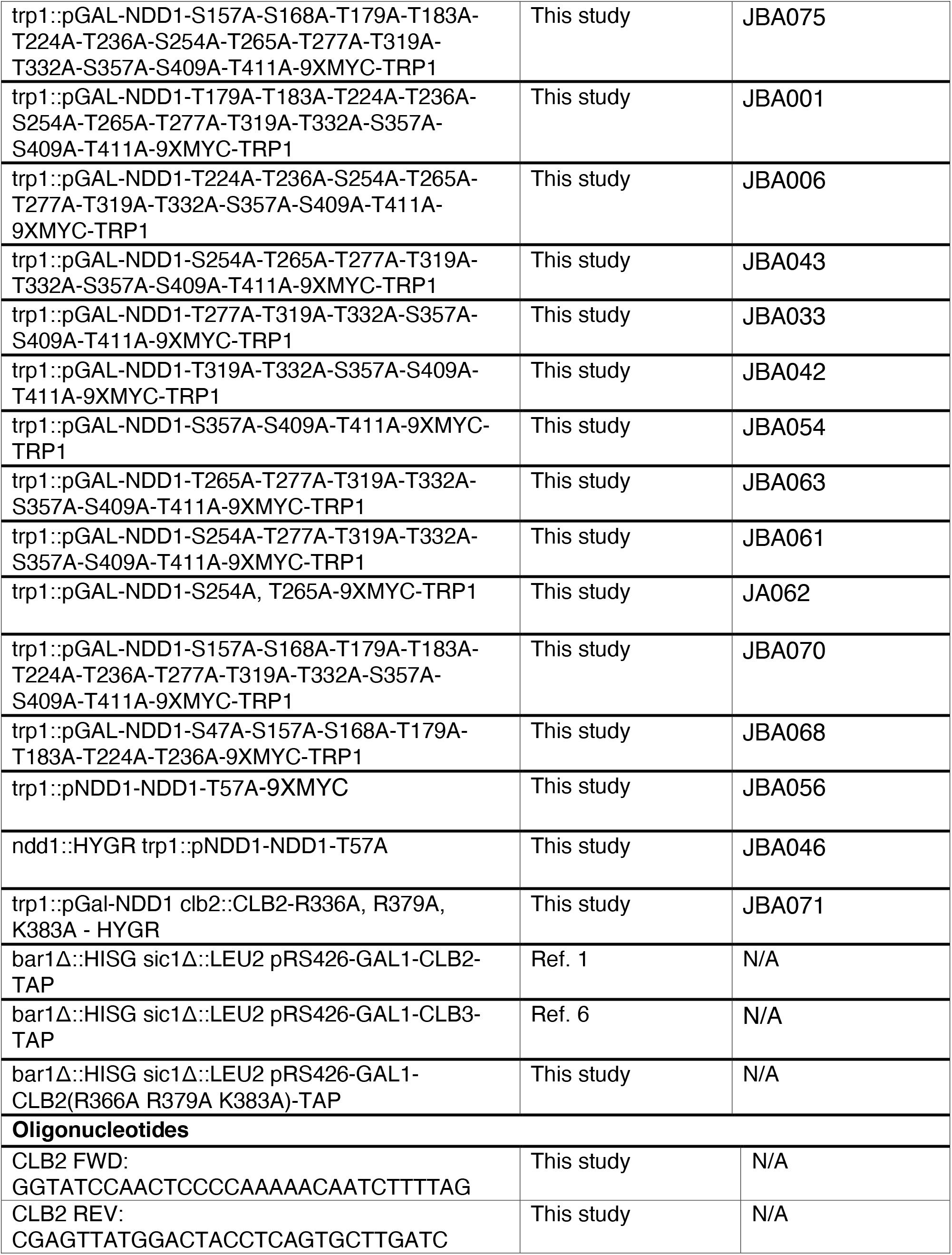
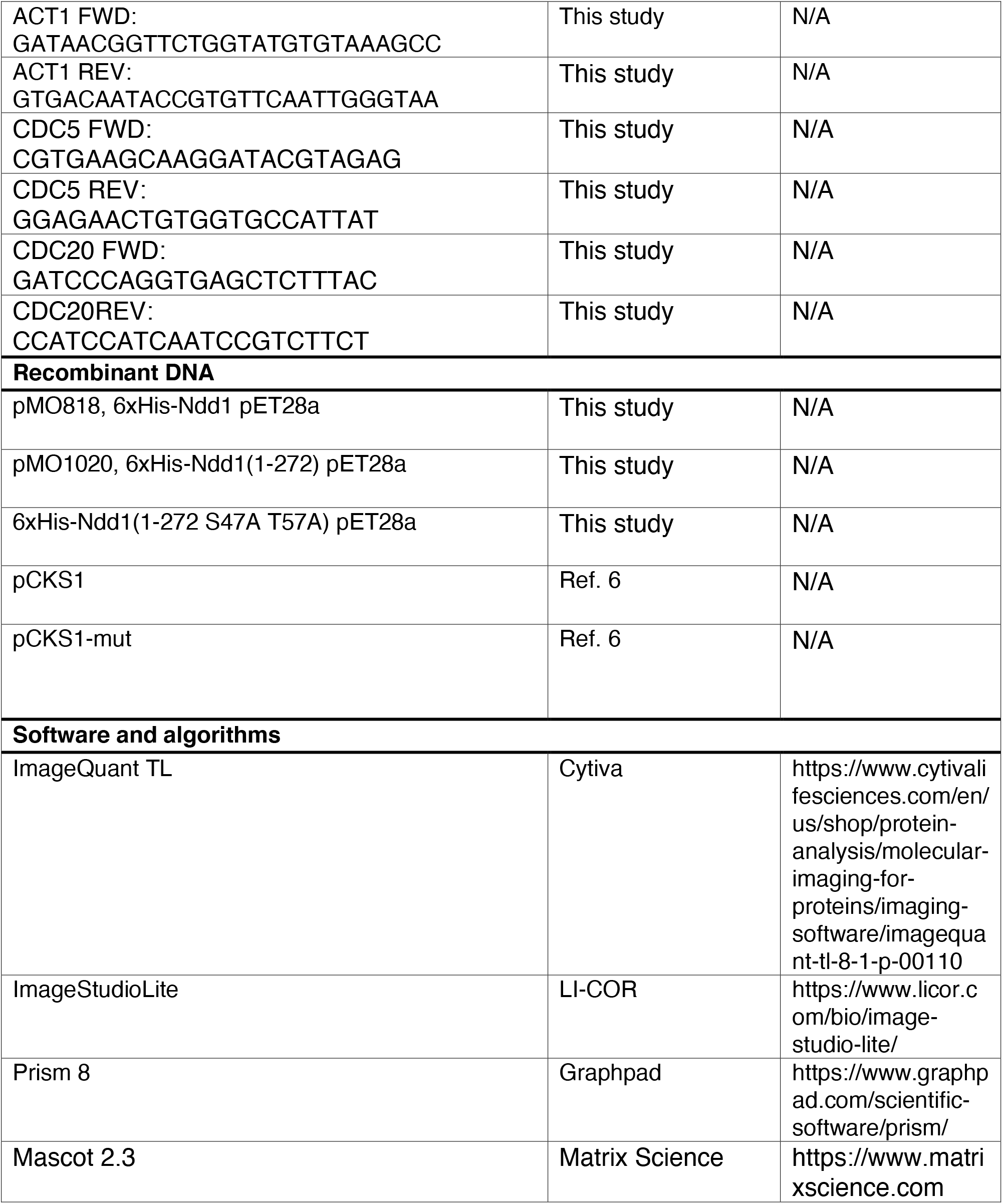

## CONTACT FOR REAGENT AND RESOURCE SHARING

Further information and requests for resources and reagents should be directed to and will be fulfilled by the Lead Contact, David Morgan.

## EXPERIMENTAL MODEL AND SUBJECT DETAILS

All strains are derivatives of W303a and listed in the Key Resources Table. All strains were constructed using PCR- and/or restriction digest-based homologous recombination. Cells were grown in Yeast Extract Peptone (YEP) containing a combination of Raffinose, Galactose, Dextrose, or a combination thereof. For all experiments, cells were grown at 30°C and progressed through a minimum of two doubling times in log phase before any experiments were performed. For all time courses, cultures reached an OD_595_ between 0.15 and 0.3 before synchronization. Cells were synchronized in G1 using 1.5 µg/ml alpha-factor for 3 h at 30°C and released by washing three times in fresh media. Cells were arrested in mitosis by addition of 15 µg/ml nocodazole for 3 h at 30°C.

## METHOD DETAILS

### Analysis of Gene Expression

Total RNA was prepared by hot-acid phenol chloroform extraction. 5% of total RNA was DNase-treated in 50 µl reactions with TURBO DNase (Thermo Fisher Scientific, Cat# AM1907) for 30 min at 37°C. 1 µl was added to a 9 µl Luna^®^ Universal One-Step RT-qPCR reaction (NEB, Cat# E3005L). RT-qPCR was carried out to manufacturer specifications in a CFX96 Touch Real Time PCR Machine (Bio-Rad). *CLB2*, *CDC5, and CDC20* expression were normalized to *ACT1* gene expression. Primers are listed in the Key Resources Table.

### Western Blotting

Ndd1 stability was analyzed in yeast strains expressing C-terminally Myc-tagged Ndd1 under the control of the *GAL1-10* promoter. *NDD1* expression was induced during mitotic synchronization by the addition of galactose to a final concentration of 2% in YEP-Raffinose. Cycloheximide was added to a final concentration of 250 µg/ml to halt translation, and 1 ml samples were harvested every 10 min to measure Ndd1 levels by Western blotting. Half-life curves were generated by fitting the data to a single exponential decay model in Prism.

For Western blotting, yeast lysates were prepared by bead-beating cells in urea lysis buffer (20 mM Tris-HCl 7.4, 7 M Urea, 2 M Thiourea, 65 mM CHAPS, 10 mM DTT). Lysates were separated using SDS-PAGE and transferred in Tris/Glycine Buffer to a 0.45 micron Nitrocellulose membrane (GE-Healthcare Life Sciences). Blots were probed with the following antibodies diluted 1:5000 in TBS-T containing 5% nonfat dry milk: mouse anti-Myc, Mouse anti-HA, Rabbit anti-Mouse IgG (H+L) Alexa Fluor 680, or 800CW Donkey anti-Mouse IgG GAPDH Monoclonal Antibody was diluted 1:1500 in TBS-T containing 5% nonfat dry milk. Blots were imaged on an Odyssey Fc imager (LI-COR) at 700 nm for 10 min.

### Analysis of Ndd1 phosphorylation

6xHis-tagged Ndd1 and Ndd1(1-272) were expressed from pET28a in *E. coli* BL21RP cells at 30°C using 0.3 mM IPTG. 6xHis-Ndd1 was purified using immobilized nickel affinity chromatography and eluted with imidazole. Clb3-and Clb2-Cdk1 complexes were purified from *S. cerevisiae* lysates using TAP-tagged cyclins as described previously^1, 43^. Cks1 was expressed in *E. coli* BL21RP and purified as described^44^.

For multisite phosphorylation analysis of Ndd1 and mutants, phosphorylation reactions were supplemented with [γ-^32^P]-ATP (Hartmann Analytic). Reactions were stopped at 5, 15, 30 and 60 min by pipetting an aliquot into SDS-PAGE sample buffer supplemented with 1 mM MnCl_2_. Reactions with full-length Ndd1 were separated using Mn-Phos-tag SDS-PAGE with 8% acrylamide and 25 µM Phos-tag (Wako Chemicals). For Ndd1(1-272), SDS-PAGE with 8% acrylamide and 50 µM Phos-tag was used. ^32^P phosphorylation signals were detected using an Amersham Typhoon 5 Biomolecular Imager (GE Healthcare Life Sciences) and quantified using ImageQuant TL (Amersham Biosciences). All kinase assays were performed in at least two replicate experiments.

For mass spectrometry analysis, kinase reactions were carried out at room temperature in 50 mM Hepes-KOH, pH 7.4, 150 mM NaCl, 5 mM MgCl_2_, 20 mM imidazole, 2% glycerol, 0.2 mg/ml BSA, and 500 µM ATP. The concentration of Ndd1 was 1 µM and of Cks1 was 500 nM. To quantitatively compare the phosphorylation of Ndd1 at different sites in different conditions, the reactions were supplemented with either normal isotopic ATP ([^16^O]ATP) or heavy ATP ([^18^O]ATP) (Cambridge Isotope Laboratories). To compare Ndd1 phosphorylation at low initial substrate turnover and high turnover, Ndd1 was phosphorylated with 0.4 nM Clb2-Cdk1 for 4 min for low turnover, or 2 nM Clb2-Cdk1 for 40 min for high turnover. For comparison of phosphorylation by Clb2- and Clb3-Cdk1, 0.4 nM Cdk1 complex was used and the reactions were stopped at 20 min. The reactions were stopped with SDS loading buffer, and aliquots from different reactions were mixed together in a 1:1 ratio and analyzed by SDS-PAGE. The gel was stained with Colloidal Coomassie G-250 and the Ndd1 band was excised. In-gel digestion was performed using trypsin/P (20 ng/µl), and peptides were purified using C18 StageTips. Peptides were separated by an Agilent 1200 series nanoflow system (Agilent Technologies) and analyzed using an LTQ Orbitrap classic mass spectrometer (Thermo Electron) equipped with a nanoelectrospray ion source (Proxeon). Mascot 2.3 (Matrix Science) was used to identify the peptides. Two independent experiments were performed.

## Acknowledgements

We thank members of the Morgan and Loog laboratories for discussions and comments on the manuscript. This work was supported by an HHMI Gilliam Graduate Fellowship (to J.B.A), a grant from the National Institute of General Medical Sciences (R35-GM118053, to D.O.M.), ERC Consolidator Grant 649124 (to M.L.), Centre of Excellence for Molecular Cell Technologies TK143 (to M.L.), and Estonian Science Agency grant PRG550 (to M.L.).

## Author contributions

J.B.A. conceived the project, performed experiments and analyzed results, with assistance from C.R.C. and guidance from D.O.M. and M.L. M.Ö., I.F., and M.L. performed Ndd1 phosphorylation analyses in vitro. J.B.A. and D.O.M. wrote the paper with assistance from all other authors.

## Supplemental Information

**Figure S1.**
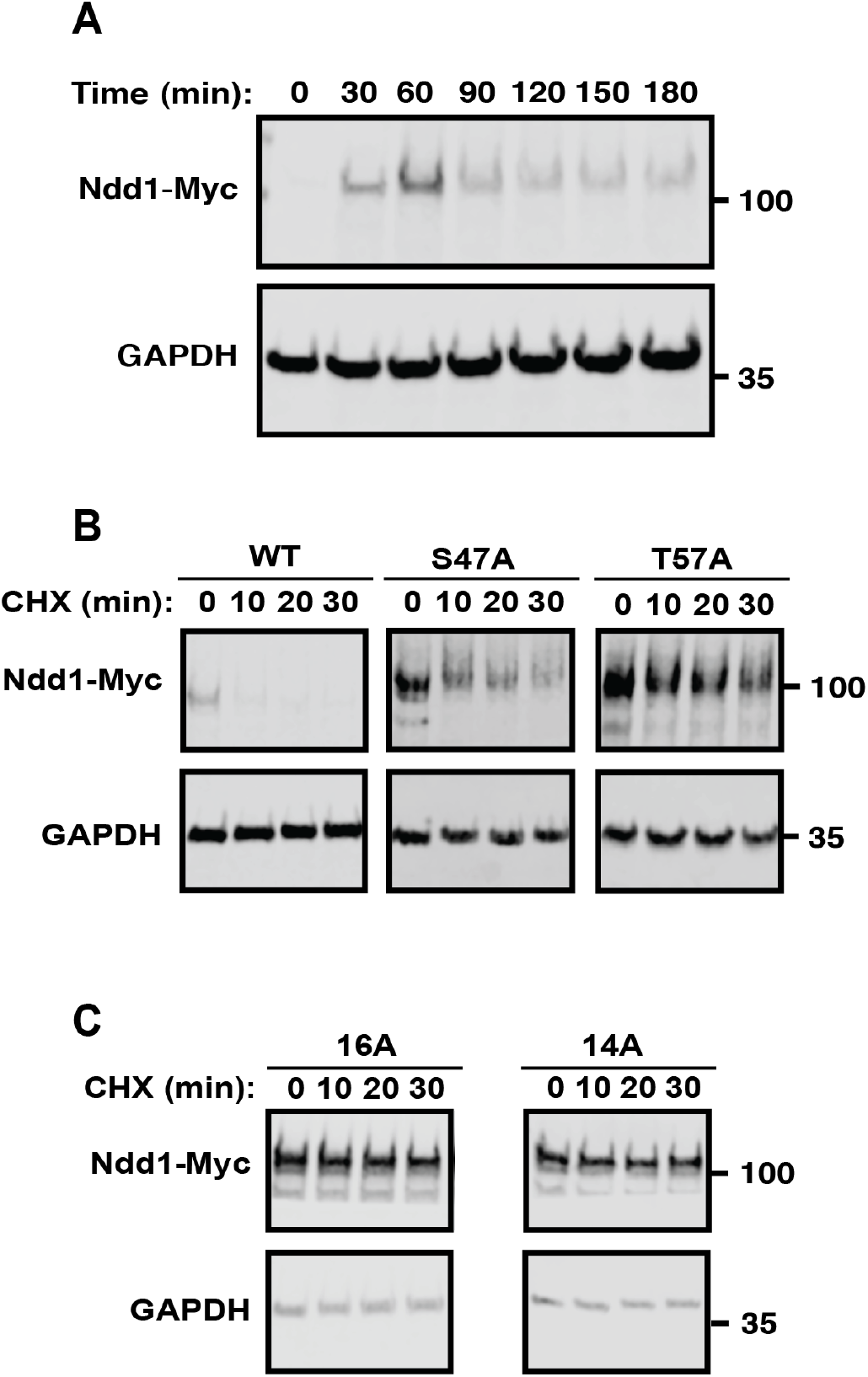
Ndd1 levels and stability in a mitotic arrest. (A) Cells were released from a G1 arrest into nocodazole. Myc-tagged Ndd1 and GAPDH expression were monitored by immunoblot at the indicated times. (B) Representative immunoblots of data quantified in Figure 2D. (C) Representative immunoblots of data quantified in Figure 2E.

**Figure S2.**
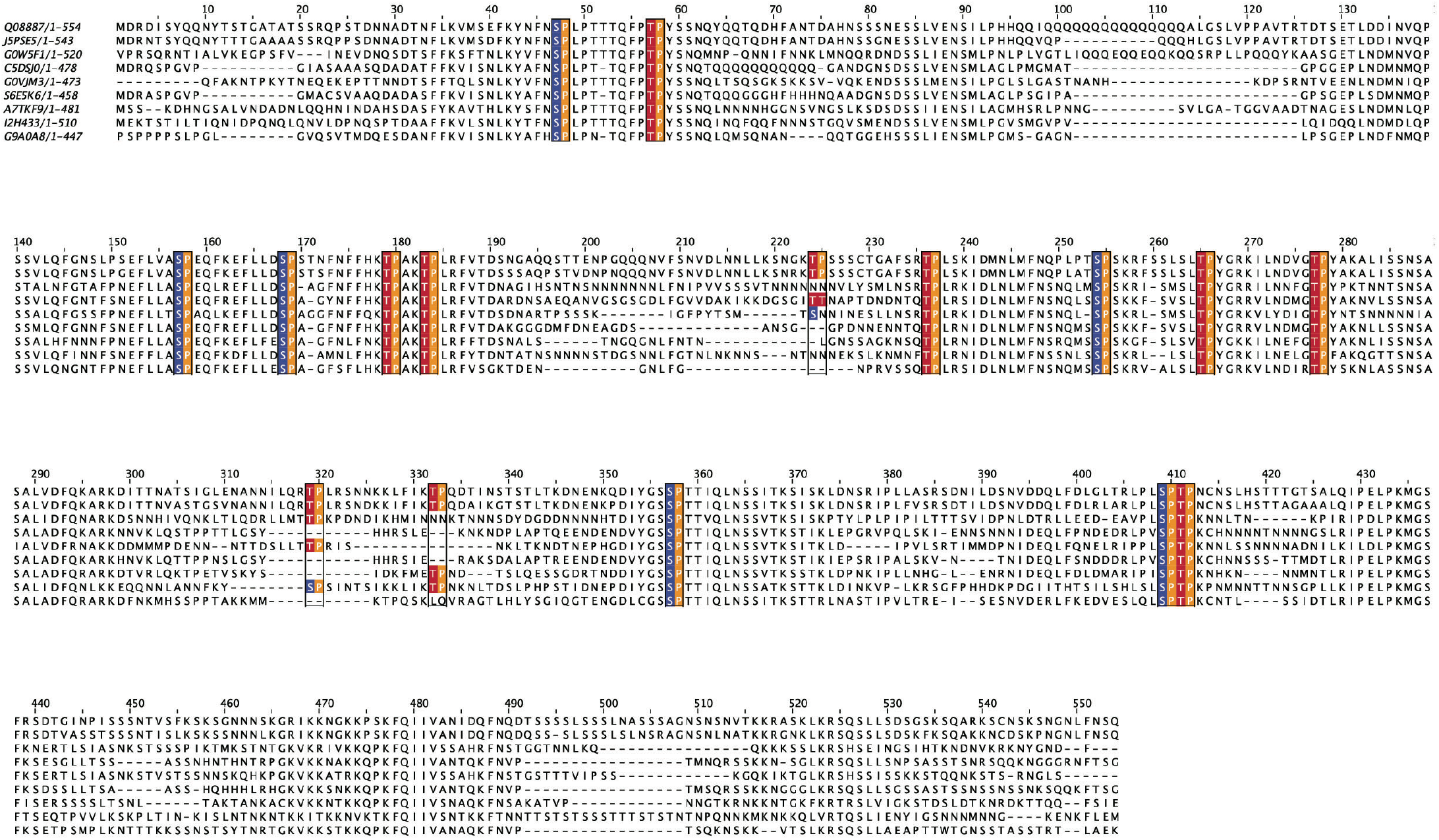
Sequence alignment of Ndd1 orthologs from multiple yeast species. The Ndd1 primary sequence from *Saccharomyces cerevisiae* was aligned to other budding yeasts using ProViz (http://slim.icr.ac.uk/proviz/). Ndd1 contains multiple sets of predicted Cdk1 sites (highlighted), spaced 10-12 resides apart. The Ndd1 sequences in the alignment are from the following species (Uniprac ID): *Saccharomyces cerevisiae* (Q08887), *Saccharomyces kudriavzevii* (J5PSE5), *Naumovozyma dairenensis* (G0W5F1), *Zygosaccharomyces rouxii* (C5DSJ0), *Naumovozyma castellii* (G0VJM3), *Zygosaccharomyces bailii* (S6E5K6), *Vanderwaltozyma polyspora* (A7TKF9), *Tetrapisispora blattae* (I2H433), *Torulaspora delbrueckii* (G9A0A8)

**Figure S3.**
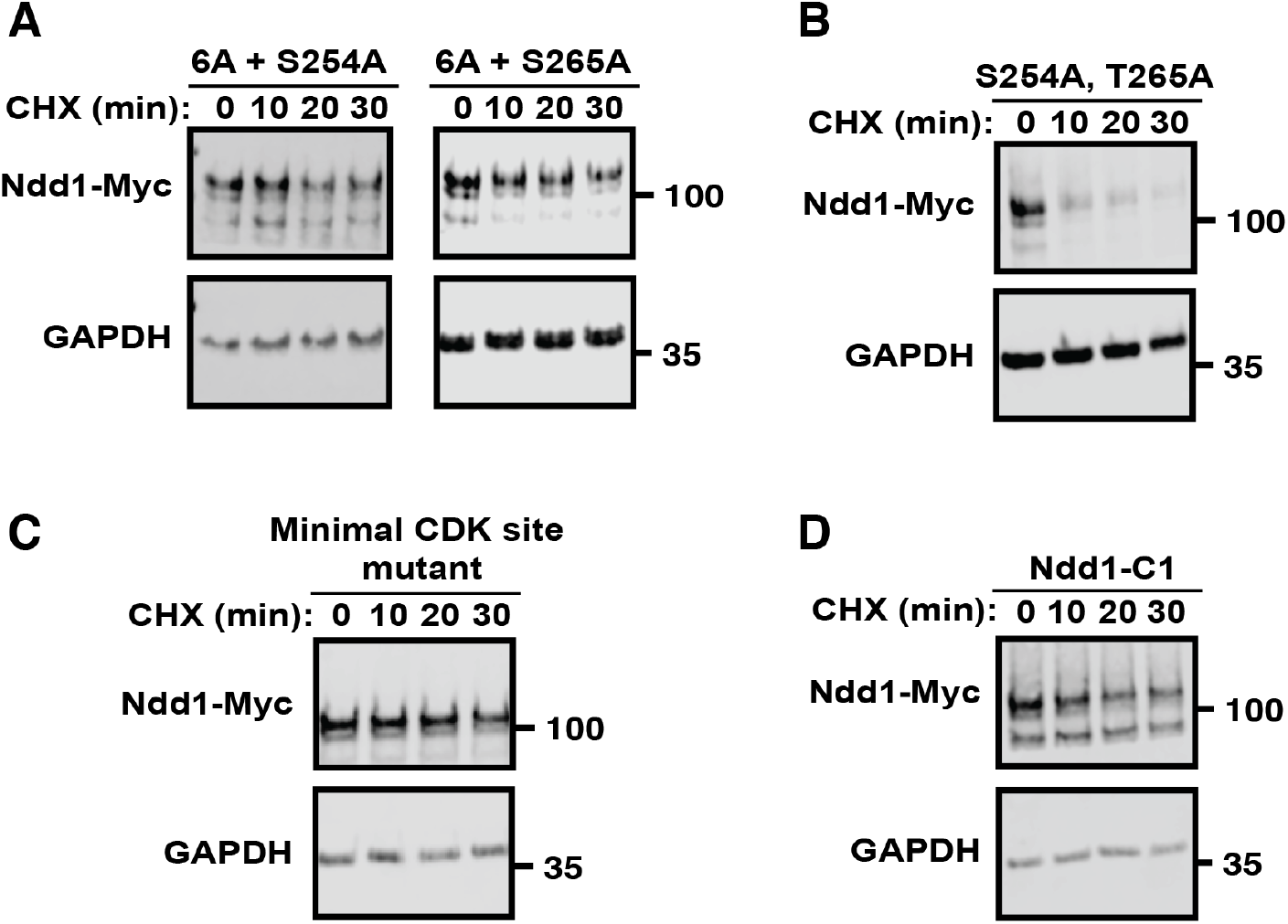
Multiple predicted Cdk1 sites promote Ndd1 destabilization. (A) Representative immunoblots of data quantified in Figure 3D. (B) Representative immunoblots of data quantified in Figure 3E. (C) Representative immunoblot of data quantified in Figure 3F. (D) Representative immunoblot of data quantified in Figure 3G.

**Figure S4.**
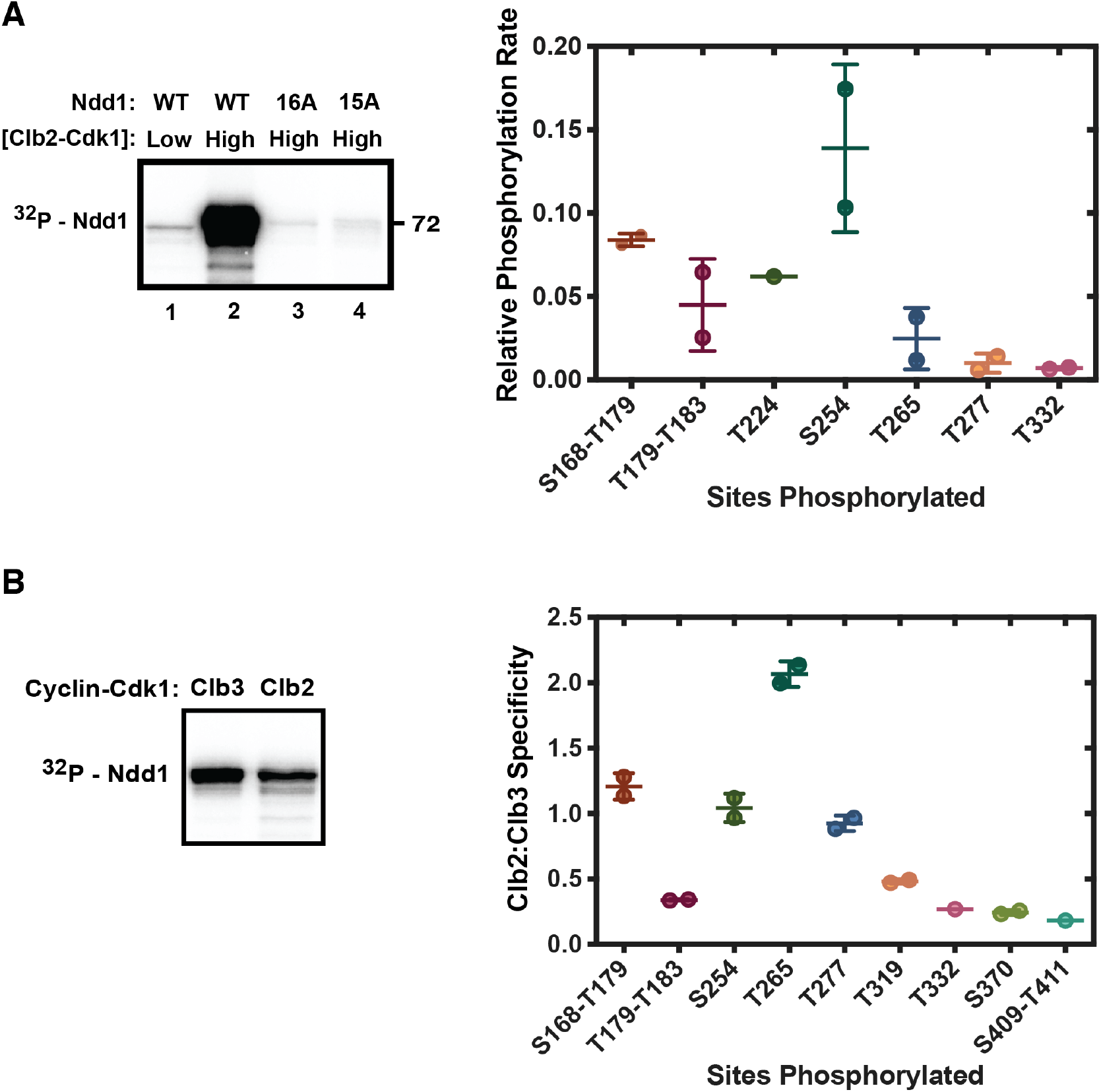
Quantitative mass spectrometry analysis of Ndd1 phosphorylation in vitro. (A) To estimate relative rates of phosphorylation at different sites in Ndd1, we compared phosphorylation after incubation of 1 µM Ndd1 at either low initial substrate turnover (4 min incubation with 400 pM Clb2-Cdk1-Cks1) or high substrate turnover (40 min incubation with 2 nM Clb2-Cdk1-Cks1). We reasoned that most sites were close to saturation at the high kinase concentration. Thus, the fraction of this maximal phosphorylation at a low (limiting) kinase concentration should provide an estimate of the relative intrinsic rate of phosphorylation at that site. In a pilot experiment (left panel), wild-type (WT) Ndd1 was phosphorylated with radiolabeled ATP in kinase reactions with 400 pM Clb2-Cdk1-Cks1 for 4 min (low, lane 1) or 2 nM Clb2-Cdk1-Cks1 for 40 min (high, lanes 2-4). In lanes 3 and 4, the high kinase concentration was used with mutant Ndd1 lacking all predicted Cdk1 sites (Ndd1-16A) or a version lacking all predicted Cdk1 sites except T57 (Ndd1-15A). Reaction products were analyzed by SDS-PAGE and autoradiography. In a parallel mass spectrometry analysis (right panel), reactions were carried out with low or high Clb2-Cdk1 concentrations as in the left panel, supplemented with either light isotopic ATP ([^16^O]ATP) or heavy isotopic ATP ([^18^O]ATP), respectively. Tryptic peptides containing the indicated phosphorylation sites were measured using mass spectrometry to quantitatively compare phosphorylation rates in the two reactions. The rate of phosphorylation at the indicated sites is expressed as the amount of phosphorylation in the low kinase reaction as a fraction of the amount of phosphorylation in the high kinase reaction. Data points (mean +/-S.D.) are from 2 independent experiments; the T224 peptide was not detected in one experiment. See Figure S5 for peptides detected in these analyses. (B) A similar experiment was performed to estimate the cyclin specificity of different sites in Ndd1. In a pilot experiment (left panel), 400 pM purified Clb3-Cdk1-Cks1 or Clb2-Cdk1-Cks1 was incubated in a kinase reaction with 1 µM Ndd1 and ^32^P-γ-ATP. After 20 min, reaction products were analyzed by SDS-PAGE and autoradiography. In a parallel mass spectrometry analysis (right panel), 400 pM Clb3- or Clb2-Cdk1 were incubated with 1 µM Ndd1 in kinase reactions supplemented with either light isotopic ATP ([^16^O]ATP) or heavy isotopic ATP ([^18^O]ATP), respectively, to quantitatively compare phosphorylation rates with the two kinases. Tryptic peptides containing the indicated phosphorylation sites were measured using mass spectrometry, and the ratio of phosphorylation by Clb2-Cdk1 to Clb3-Cdk1 is indicated. Data points (mean +/-S.D.) are from 2 independent experiments. Two peptides were not detected in one experiment, and the T224 peptide observed in panel A was not detected in either experiment here. See Figure S5 for peptides detected in these analyses.

**Figure S5.**
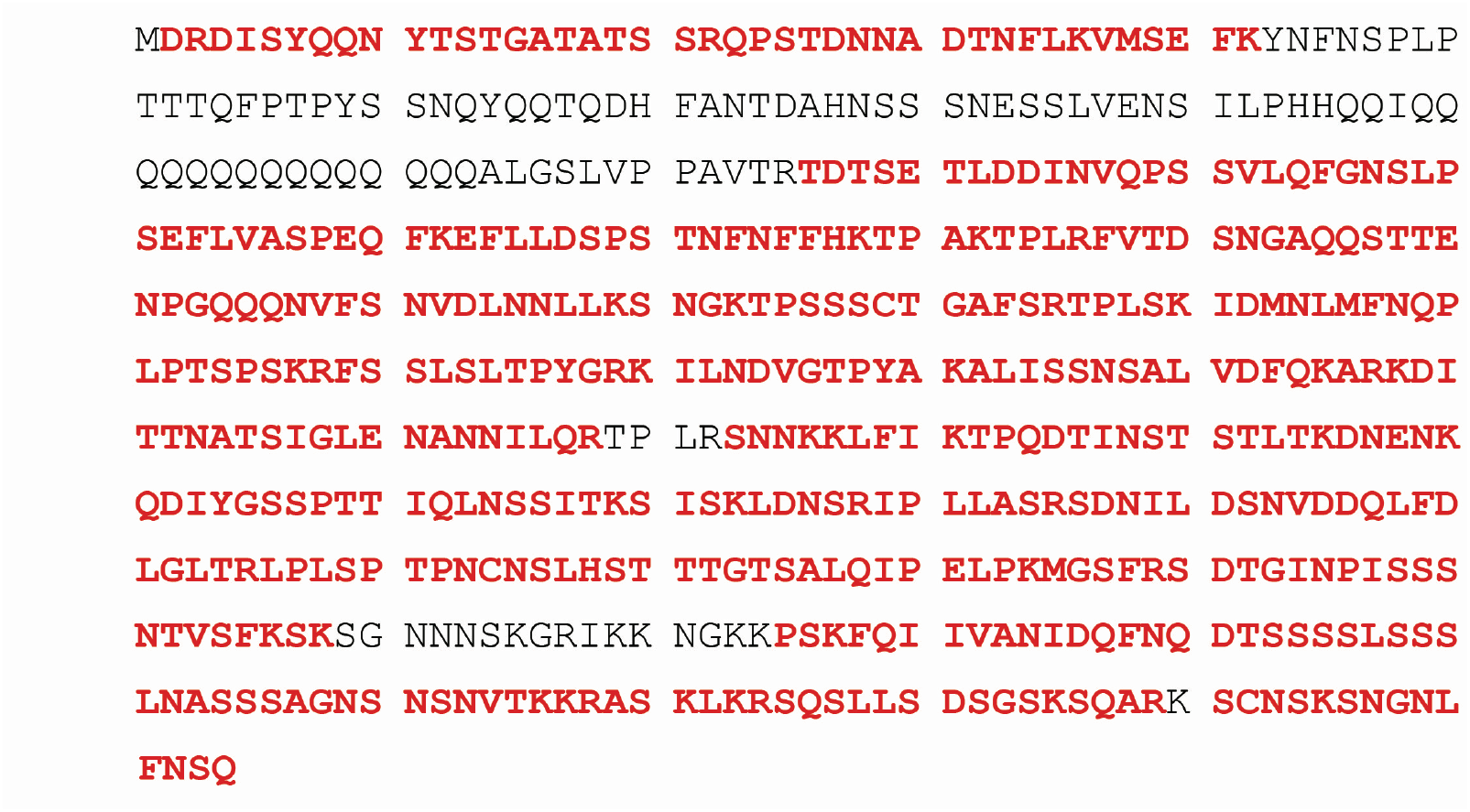
Ndd1 mass spectrometry sequence coverage. Peptides containing sequences detected by mass spectrometry in Figure S4A and B are indicated in bold red text. Sequences that were not detected are indicated in black text.

## References

1. Ubersax, J. A. et al. Targets of the cyclin-dependent kinase Cdk1. Nature 425, 859–864 (2003).

2. Holt, L. J. et al. Global analysis of Cdk1 substrate phosphorylation sites provides insights into evolution. Science 325, 1682–1686 (2009).

3. Loog, M. & Morgan, D. O. Cyclin specificity in the phosphorylation of cyclin-dependent kinase substrates. Nature 434, 104–108 (2005).

4. Swaffer, M. P., Jones, A. W., Flynn, H. R., Snijders, A. P. & Nurse, P. CDK Substrate Phosphorylation and Ordering the Cell Cycle. Cell 167, 1750–1761.e16 (2016).

5. Coudreuse, D. & Nurse, P. Driving the cell cycle with a minimal CDK control network. Nature 468, 1074–1079 (2010).

6. Kõivomägi, M. et al. Cascades of multisite phosphorylation control Sic1 destruction at the onset of S phase. Nature 480, 128–131 (2011).

7. Pirincci Ercan, D., et al. Budding yeast relies on G1 cyclin specificity to couple cell cycle progression with morphogenetic development. Sci Adv 7, eabg0007 (2021).

8. Bhaduri, S. & Pryciak, P. M. Cyclin-specific docking motifs promote phosphorylation of yeast signaling proteins by G1/S Cdk complexes. Curr. Biol. 21, 1615–1623 (2011).

9. Örd, M., Venta, R., Möll, K., Valk, E. & Loog, M. Cyclin-Specific Docking Mechanisms Reveal the Complexity of M-CDK Function in the Cell Cycle. Molecular Cell (2019). doi:10.1016/j.molcel.2019.04.026

10. Örd, M. et al. Multisite phosphorylation code of CDK. Nat Struct Mol Biol 26, 649– 658 (2019).

11. Lyons, N. A., Fonslow, B. R., Diedrich, J. K., Yates, J. R. & Morgan, D. O. Sequential primed kinases create a damage-responsive phosphodegron on Eco1. Nat Struct Mol Biol 20, 194–201 (2013).

12. Kõivomägi, M. et al. Multisite phosphorylation networks as signal processors for Cdk1. Nat Struct Mol Biol 20, 1415–1424 (2013).

13. Valk, E. et al. Multistep phosphorylation systems: tunable components of biological signaling circuits. Molecular Biology of the Cell 25, 3456–3460 (2014).

14. Ferrell, J. E. & Ha, S. H. Ultrasensitivity part II: multisite phosphorylation, stoichiometric inhibitors, and positive feedback. Trends in Biochemical Sciences 39, 556–569 (2014).

15. Wittenberg, C. & Reed, S. I. Cell cycle-dependent transcription in yeast: promoters, transcription factors, and transcriptomes. Oncogene 24, 2746–2755 (2005).

16. Spellman, P. T. et al. Comprehensive identification of cell cycle-regulated genes of the yeast Saccharomyces cerevisiae by microarray hybridization. Molecular Biology of the Cell 9, 3273–3297 (1998).

17. Kumar, R., Reynolds, D. M., Shevchenko, A., Goldstone, S. D. & Dalton, S. Forkhead transcription factors, Fkh1p and Fkh2p, collaborate with Mcm1p to control transcription required for M-phase. Curr. Biol. 10, 896–906 (2000).

18. Pic-Taylor, A., Darieva, Z., Morgan, B. A. & Sharrocks, A. D. Regulation of cell cycle-specific gene expression through cyclin-dependent kinase-mediated phosphorylation of the forkhead transcription factor Fkh2p. Molecular and Cellular Biology 24, 10036–10046 (2004).

19. Reynolds, D. et al. Recruitment of Thr 319-phosphorylated Ndd1p to the FHA domain of Fkh2p requires Clb kinase activity: a mechanism for CLB cluster gene activation. Genes Dev. 17, 1789–1802 (2003).

20. Darieva, Z. et al. Cell cycle-regulated transcription through the FHA domain of Fkh2p and the coactivator Ndd1p. Curr. Biol. 13, 1740–1745 (2003).

21. Yu, J. et al. Structural basis of human separase regulation by securin and Cdk1-cyclin B1. Nature (2021) https://doi.org/10.1038/s41586-021-03764-0.

22. Surana, U. et al. Destruction of the CDC28/CLB mitotic kinase is not required for the metaphase to anaphase transition in budding yeast. EMBO J 12, 1969–1978 (1993).

23. Linke, C. et al. A Clb/Cdk1-mediated regulation of Fkh2 synchronizes CLB expression in the budding yeast cell cycle. NPJ Syst Biol Appl 3, 7–12 (2017).

24. Palframan, W. J., Meehl, J. B., Jaspersen, S. L., Winey, M. & Murray, A. W. Anaphase inactivation of the spindle checkpoint. Science 313, 680–684 (2006).

25. Vernieri, C., Chiroli, E., Francia, V., Gross, F. & Ciliberto, A. Adaptation to the spindle checkpoint is regulated by the interplay between Cdc28/Clbs and PP2ACdc55. The Journal of Cell Biology 202, 765–778 (2013).

26. Alexandru, G., Zachariae, W., Schleiffer, A. & Nasmyth, K. Sister chromatid separation and chromosome re-duplication are regulated by different mechanisms in response to spindle damage. EMBO J 18, 2707–2721 (1999).

27. Amon, A., Tyers, M., Futcher, B. & Nasmyth, K. Mechanisms that help the yeast cell cycle clock tick: G2 cyclins transcriptionally activate G2 cyclins and repress G1 cyclins. Cell 74, 993–1007 (1993).

28. Bishop, A. C. et al. A chemical switch for inhibitor-sensitive alleles of any protein kinase. Nature 407, 395–401 (2000).

29. Cardozo, T. & Pagano, M. The SCF ubiquitin ligase: insights into a molecular machine. Nat Rev Mol Cell Biol 5, 739–751 (2004).

30. Berset, C. et al. Transferable domain in the G(1) cyclin Cln2 sufficient to switch degradation of Sic1 from the E3 ubiquitin ligase SCF(Cdc4) to SCF(Grr1). Molecular and Cellular Biology 22, 4463–4476 (2002).

31. Hsiung, Y. G. et al. F-box protein Grr1 interacts with phosphorylated targets via the cationic surface of its leucine-rich repeat. Molecular and Cellular Biology 21, 2506–2520 (2001).

32. Edenberg, E. R., Mark, K. G. & Toczyski, D. P. Ndd1 turnover by SCF(Grr1) is inhibited by the DNA damage checkpoint in Saccharomyces cerevisiae. PLoS Genet. 11, e1005162 (2015).

33. Örd, M. et al. Proline-Rich Motifs Control G2-CDK Target Phosphorylation and Priming an Anchoring Protein for Polo Kinase Localization. Cell Rep 31, 107757 (2020).

34. Kõivomägi, M. et al. Dynamics of Cdk1 substrate specificity during the cell cycle. Molecular Cell 42, 610–623 (2011).

35. McGrath, D. A. et al. Cks confers specificity to phosphorylation-dependent CDK signaling pathways. Nat Struct Mol Biol 20, 1407–1414 (2013).

36. Örd, M. & Loog, M. Detection of Multisite Phosphorylation of Intrinsically Disordered Proteins Using Phos-tag SDS-PAGE. Methods Mol. Biol. 2141, 779– 792 (2020).

37. Brandman, O. & Meyer, T. Feedback loops shape cellular signals in space and time. Science 322, 390–395 (2008).

38. Goda, T., Ishii, T., Nakajo, N., Sagata, N. & Kobayashi, H. The RRASK motif in Xenopus cyclin B2 is required for the substrate recognition of Cdc25C by the cyclin B-Cdc2 complex. J. Biol. Chem. 278, 19032–19037 (2003).

39. Blondel, M. et al. Degradation of Hof1 by SCF(Grr1) is important for actomyosin contraction during cytokinesis in yeast. EMBO J 24, 1440–1452 (2005).

40. Quilis, I. & Igual, J. C. A comparative study of the degradation of yeast cyclins Cln1 and Cln2. FEBS Open Bio 7, 74–87 (2017).

41. Salama, S. R., Hendricks, K. B. & Thorner, J. G1 cyclin degradation: the PEST motif of yeast Cln2 is necessary, but not sufficient, for rapid protein turnover. Molecular and Cellular Biology 14, 7953–7966 (1994).

42. Darieva, Z. et al. Polo kinase controls cell-cycle-dependent transcription by targeting a coactivator protein. Nature 444, 494–498 (2006).

43. Puig, O. et al. The tandem affinity purification (TAP) method: a general procedure of protein complex purification. Methods 24, 218–229 (2001).

44. Reynard, G. J., Reynolds, W., Verma, R. & Deshaies, R. J. Cks1 is required for G(1) cyclin-cyclin-dependent kinase activity in budding yeast. Molecular and Cellular Biology 20, 5858–5864 (2000).

